# Biomimetic model for the identification of distinctive microenvironmental factors in glioblastoma radiosensitivity

**DOI:** 10.64898/2026.01.14.699525

**Authors:** Ryan Yao, Isabella Rivera, Jorge Maldonado, Catherine A Best-Popescu, Hui Xu, Jann N Sarkaria, Jingxiang Wang, Yi Lu, Kim A Selting, Brendan A.C. Harley, Sara Pedrón-Haba

## Abstract

Radiation therapy (RT) has long been included in the treatment of glioblastoma (GBM). However, radioresistance in cancer cells as well as toxicity in normal tissues are major obstacles to clinical efficacy. Improved understanding of the mechanisms of tumor microenvironment-induced radioresistance during and after radiation therapy can provide fundamental insights to improve clinical outcomes in GBM. Here, using three-dimensional engineered hydrogel models in vitro, we report the influence of extracellular matrix, hypoxia, and adjacent neuronal cells in radiotherapeutic sensitivity. We find that mechanical cues and oxygen availability regulate cellular response to radiation, with softer matrices allowing for more DNA damage. Hyaluronan fragments from the extracellular matrix also modulate rapid metabolic response to radiation, especially in hypoxic environments. We show that neuronal networks influence tumor metabolic activity and the inflammatory response. Overall, we demonstrate here that alternative radiation strategies, such as low dose rate radiation therapy and microenvironmental regulation, have the potential to be more effective in a specific subset of radiosensitive GBM tumors.

## Introduction

Glioblastoma (GBM) is a common primary malignant brain tumor in adults with a poor prognosis (5-year survival rate less than 7%) despite treatment. Patients face limited treatment options, due to fast infiltration, tumor heterogeneity, and immune suppression which leads to relapse. Maximal resection of the tumor mass is the first treatment option, and recently, increasing reports suggest supramarginal resection as a strategy to increase overall survival. However, challenges exist in the establishment of surgical margins without producing additional neurological deficits (1). Radiation therapy is also a standard of care for GBM patients, increasing survival when compared to surgery alone, and often delivered with concomitant temozolomide (TMZ) treatment. The main purpose of radiation therapy is to achieve local tumor control while minimizing toxicity in adjacent normal tissue, thus improving therapeutic ratio. A better understanding of the biologic response to radiation of GBM tumors and the normal parenchyma may help inform treatments that enhance the therapeutic ratio (2). Methods to improve response to therapy while minimizing normal tissue toxicity include image-guided radiation therapy for a more precise delivery of RT, new dose rates and protocols that exploit differences in radiation response between tumor and normal tissue, and the co-administration of therapeutic agents that modify the tumor microenvironment or increase the tumor radiosensitivity (3). Despite efforts to provide more precise RT treatments, radiation-related toxicity in the brain continues to be problematic leading to neurological deficits in patients with long survival.

Ionizing radiation induces significant DNA damage leading ultimately to tumor cell death. Among these lesions, DNA double strand breaks (DSBs) are difficult to repair and thus most likely to lead to replicative sterility. The main mechanisms of repair includes protein kinases, and their inhibition has led to increased radiosensitivity in head and neck cancer (4). The histone H2AX is phosphorylated to γ-H2AX in response to DSBs and γ-H2AX foci can be used as a biomarker for irreparable damage. γ-H2AX can be visualized by immunofluorescence through primary and secondary antibodies (5). The center of this cytotoxic process is the generation of reactive oxygen species (ROS); RT can damage mitochondrial function, leading to sustained production of ROS. Because of the importance of reactive oxygen species in DNA damage after radiation, hypoxia and re-oxygenation are essential factors in RT sensitivity (6). The understanding of the cellular and molecular mechanism of neoplastic mitochondria and metabolism to enhance radioresistance will allow for the development of strategies that target cellular bioenergetics that increase radiosensitivity.

Alternative radiation regimens and rates have been explored to increase the therapeutic ratio, taking advantage of the differences in cellular processes including DNA repair between normal and tumor tissue. Recent data show that a single-fraction brain irradiation delivered by ultra-high dose rate radiotherapy (>40 Gy/sec) causes the FLASH effect (FLASH-RT) and does not elicit the neurocognitive deficits in mice that are seen with conventional dose rate RT (∼6 Gy/min) (3,7). Low dose rate (<1 Gy/min) has also been explored as an alternative to conventional rates. This strategy takes advantage of the sublethal damage repair of normal tissue, associated with decreased release of transforming growth factor-ß, and increasing the therapeutic ratio through improved tumor control as a result of low-dose hypersensitivity (8,9). This process allows for improved restoration of function in normal tissues while delivering a therapeutic dose to the tumor.

Matrix stiffness can strongly influence tumor response to radiation therapy. Tumors show a heterogenous distribution of mechanical properties, that dynamically changes during tumor progression due to continuous degradation and secretion of different extracellular matrix (ECM) components. In addition, RT has been associated with changes in tumor architecture and stiffness that affect tumor cell sensitivity as a result. Moreover, previous studies show that irradiated glioma tumors reprogram glucose and serine metabolism to promote antioxidant responses that hinder the cytotoxic effects of radiation therapy and promote unsaturated fatty acids accumulation that prevents immune cell recruitment (10–12). The discovery of connections between radiation and components of the tumor microenvironment can improve clinical outcomes of RT for glioblastoma patients (13).

Here we describe the use of a gelatin hydrogel system to evaluate the role of the extracellular matrix (stiffness, hyaluronic acid content), gelation-induced hypoxia, and inclusion of additional cell populations from the GBM microenvironment on radiation resistance and damage. We have previously described the development of gelatin hydrogels to study processes of glioblastoma proliferation, invasion, and drug resistance (14–18) as well as radiation induced damage to vascular cells embedded within the matrix (19). When compared to traditional 2D in vitro cell culture, 3D platforms provide an avenue to capture structural, biophysical, and biochemical features of the native tumor microenvironment that are inaccessible in vivo as well as interrogate therapeutic responses to dissect mechanisms associated with GBM-radiobiology (20). We use a gelation hydrogel system to tune multiple microenvironmental factors to interrogate how the local tumor microenvironment (TME) and surrounding stroma respond to radiation. We report stromal characteristics that distinguish GBM cell response to radiotherapy. This will lead to the capacity to predict more precise radiotherapy treatments, reducing side effects that may induce cognitive impairment.

## Materials and Methods

### Hydrogel fabrication

Normoxic hydrogels were prepared with gelatin methacrylamide (GelMA). Prepolymer solution was made with GelMA (5 wt.%) and 0.1 wt.% LAP (lithium phenyl-2,4,6 trimethylbenzoylphosphinate) photoinitiator. The solution was pipetted into Teflon molds (5mm diameter and 1.5mm thickness) and exposed to 10 mW cm^-2^ UV light for 40 seconds. Hypoxic hydrogels were prepared with gelatin ferulic acid (GelFA) (21). Laccase from *Streptomyces coelicolor* was expressed heterogeneously in *Escherichia coli* BL21(DE3) and purified using previously reported methods (22). The prepolymer solution was made with GelFA (3 wt.%) and 0.06 mM laccase enzyme (21). The solution was pipetted into Teflon molds and allowed to sit for 15 min to fully polymerize before transferring to a well plate with neuronal media. Patient-derived glioblastoma lines (23) (GBM6, GBM8, GBM39, GBM12, GBM34, GBM44, and U87) were cultured in these hydrogels at a concentration of 5 million cells per ml, while primary neurons were cultured at 2 million cells per ml (24).

### Metabolic activity

The total metabolic activity of cell-containing hydrogels was measured on day 14 of hydrogel encapsulation (day 7 post low or conventional dose rate of radiation at 8 Gy). Metabolic activity was analyzed using a dimethylthiazol-diphenyltetrazolium bromide assay (MTT; Molecular Probes) following manufacturer’s instructions. Briefly, the culture media of each sample was replaced with MTT-containing media and incubated for 4 h. The solution was replaced with dimethyl sulfoxide (DMSO; Sigma-Aldrich) and set overnight. Metabolic activity of samples was measured via absorbance at 540 nm using a microplate reader (Synergy HT, Biotek). We acknowledge that MTT test may underestimate the inhibition potential of ionizing radiation, due to radiation-induced mitochondrial biogenesis and hyperactivation, however, it provides accurate trends and results are corroborated with the quantification γ-H2AX foci (25).

### Mechanical properties

Prepolymer solutions for GelMA (5 wt.% and 6 wt.%) and GelFA (0.04 mM and 0.06 mM laccase) were prepared, pipetted into 8 mm diameter Teflon molds, and polymerized as previously described. After 24 hours, half of the samples were subjected to 8 Gy of radiation. Storage and loss moduli for the hydrogels were collected with a TA DHR-3 Rheometer using an 8 mm cross-hatched plate. Samples were tested at 25°C and 3% strain over a logarithmic range of 100-0.01 rad s^-1^. Data points where the phase offset exceeded 90 degrees were excluded.

### Seahorse metabolism

Mitochondria stress test assay and ATP production rate assay were performed using a Seahorse XFe96 analyzer (Agilent) under hypoxia or normoxia conditions (26). Hydrogels were prepared in small molds to hold 20 μl and polymerized as previously described. GBM cells within hydrogels were seeded at a density of 1 × 10^5^ cells/well in Seahorse XFe96 Cell Culture microplates (Agilent, 103794-100). On the day of the assay the growth medium was replaced with assay medium (Agilent) as instructed. Cells were incubated in a non-CO_2_ incubator for 1 h before initiation of the test. The assay medium was Seahorse XF base medium supplemented with 10 mM glucose, 1 mM pyruvate and 2 mM glutamine. For the mitochondria stress test assay, the final concentrations of specific chemicals were optimized to be oligomycin (Sigma-Aldrich) at 20 μM, carbonyl cyanide-4-(trifluoromethoxy) phenylhydrazone (FCCP) (Sigma-Aldrich) at 7.5 μM, and antimycin A/rotenone (Sigma-Aldrich) at 5 μM/10 μM. The ATP production rate assay was performed with specific chemicals of 20 μM of oligomycin and antimycin A/ Rotenone at 5 μM/10 μM. The assay protocol was modified from the standard protocol by inserting an extra step after each injection of the reagents. The extra step includes 2 cycles of 3-min mix and 7-min wait. For assays under hypoxia conditions, the WAVE software was in hypoxia mode, and the assays were performed inside a hypoxia chamber following the suggested protocol. The oxygen level was set to be 3%.

### Hyaluronic acid (HA) and neuronal imaging

GBM hydrogels were fixed with 4% Image-iT paraformaldehyde in 0.1M Phosphate Buffer and permeabilized in a 0.5% Tween solution followed by three washes in 0.1% Tween solution (PBST). Samples were blocked in a 2% bovine serum albumin (BSA) solution followed by incubation with biotinylated HABP (Hyaluronic Acid Binding Protein, 1:100) and 2% BSA (Bovine Serum Albumin) at 4 °C, then Streptavidin-488 (1:500). Samples were incubated in a 1:500 dilution of DAPI in PBS for counterstaining. Neuronal and GBM hydrogels were fixed with 4% Image-iT paraformaldehyde in 0.1M Phosphate Buffer. Samples were permeabilized and blocked as described above. Neuronal samples were incubated overnight in a solution containing β-III-tubulin and 2% BSA (1:200) at 4 °C. GBM samples were incubated overnight in a solution containing Ki-67 and 2% BSA (1:500) at 4 °C. Following washes in PBST, AlexaFluor-488 conjugate was diluted in 2% BSA (1:500) and added. Samples were then incubated in a 1:300 dilution of DAPI in PBS and washed in PBST. Samples were stored in PBS until imaging (Zeiss LSM880).

### TGF-β1 analysis

ELISA was performed to identify the presence of the TGF-β biomarker in media samples collected after RT (R&D Systems #DY240), following manufacturer’s instructions. Briefly, a 96 well-plate was coated with the capture antibody; after washing (0.05% Tween 20 in PBS), the plate was blocked at room temperature for one hour. The 100 μl aliquots of media samples were activated through the addition of 20 μl 1N HCl and 20 μl of 1.2 N NaOH/0.5M HEPES. The samples’ absorbances were read with a microplate reader at 450 nm with wavelength correction at 540 nm.

### X-ray and Spatial Light Interference Microscopy (SLIM)

A W-transmission mini-Xray unit operating at 50kVp/80μA with a 2 mm aperture was developed and mounted by Dan Ionescu at University Cincinnati Ohio. The X-ray spectrum was evaluated using a CdTe spectrometer and the half-value layer (HVL) determined using aluminum foils. Output was measured using a TG61 chamber with low-energy calibration and verified using 6MV calibration with energy- and attenuation-corrected radiochromic film. Spatial Light Interference Microscopy takes advantage of the fact that optical phase delay accumulated through a live cell is linearly proportional to the cell’s non-aqueous content, enabling direct extraction of cellular dry mass from quantitative phase measurements (27). Shielding was affixed to the microscope incubator. Time-resolved cellular morphology, as well as cellular and nuclear dry-mass of incubated cells, was observed during two 20 Gy fractions delivered over 48 hours. The system is composed of a SLIM module (Cell Vista SLIM pro, Phi Optics, Inc.) attached to a commercial phase contrast microscope (axio Observer Zi, Zeiss). The SLIM module contains a 4f optical relay and a spatial light modulator (SLM) that encodes controlled phase shifts. A scientific CMOS camera records a series of intensity images that are computationally combined to reconstruct quantitative phase maps of the field of view, following the original SLIM instrumentation framework (28). From these phase maps, we obtain quantitative information about GBM and neuronal cellular density, spatial distribution, connectivity, migration, and transformation frequency.

### Analysis of γ-H2AX foci

Cells were grown in GelMA hydrogels and irradiated at 8 Gy, at low and conventional dose rates, and then further incubated for 1 h. Afterwards, the cells were fixed with PBS containing 4% (w/v) paraformaldehyde and permeabilized with 0.1% (w/v) Triton™ X-100 in PBS at room temperature. The cells were then treated with blocking buffer 0.1% (w/v) FBS, incubated with an anti-γ-H2AX antibody (Millipore, MA, USA), and then with Alexa Fluor-488 (A21425; Invitrogen) at room temperature. The nuclei of the labeled cells were counterstained with 40,6-diamidino-2-phenylindole (DAPI, D1306 Thermofisher), and samples were visualized under an LSM880 microscope (Carl Zeiss, Oberkochen, Germany). Cells displaying foci were counted as positive regarded as having DNA damage, 300 cells per condition were analyzed and 3 technical repetitions.

### Irradiation of cells in 3D culture

Plates were positioned on 1.5 cm of solid water through which the radiation beam was delivered. From beneath the solid water (used to create dose build up), a 6 MV photon beam with flattening filter was used to deliver 8 Gy of radiation with a Varian® TrueBeam® linear accelerator. Radiation was delivered at standard (6 Gy/min) or low (0.4 Gy/min) dose rates.

### Statistical analysis

All analyses were performed using a one-way analysis of variance (ANOVA) followed by Tukey’s HSD post-hoc test. Significance level was set at p < 0.05 or p < 0.01. At least n = 3 samples were examined. Error was reported in the figures as the standard deviation unless otherwise noted.

## Results and discussion

### Tumor microenvironment characterization in a 3D in vitro model

Bioengineered models have shown immense potential in recreating human tumor physiology to better understand tumor development and screening of therapeutic targets. We have fabricated and characterized biomaterial-based glioma models extensively and are able to control the biophysical properties of the extracellular matrix (ECM), cell composition, and oxygen concentration. These factors have an influence in tumor growth, invasion, and drug resistance (14–16,18). As the main component of the brain ECM, hyaluronic acid has emerged as an essential component of the glioma cell metabolism (29). The platform used for this study consists of a gelatin-based hydrogel, that can control oxygen concentration by varying the crosslinking method and allows for the monitoring of cell behavior and physical properties. The Image-IT reagent (I14833, ThermoFisher) was used to corroborate oxygen concentrations under 5%. We observe that cell metabolic activity decreases under hypoxia-inducible hydrogels, even in the presence of HA metabolites (**Figure 1A-B**), and that there is a decrease in metabolic activity in the presence of high molecular weight HA fragments. It has been reported that glioma cells show more invasive behavior under hypoxic conditions (30) and that low molecular weight HA increases mitochondrial respiration (29), suggesting that when hypoxic, glioma cells can switch from aerobic glycolysis into oxidative phosphorylation, by the secretion of selected metabolites.

**Figure 1.**
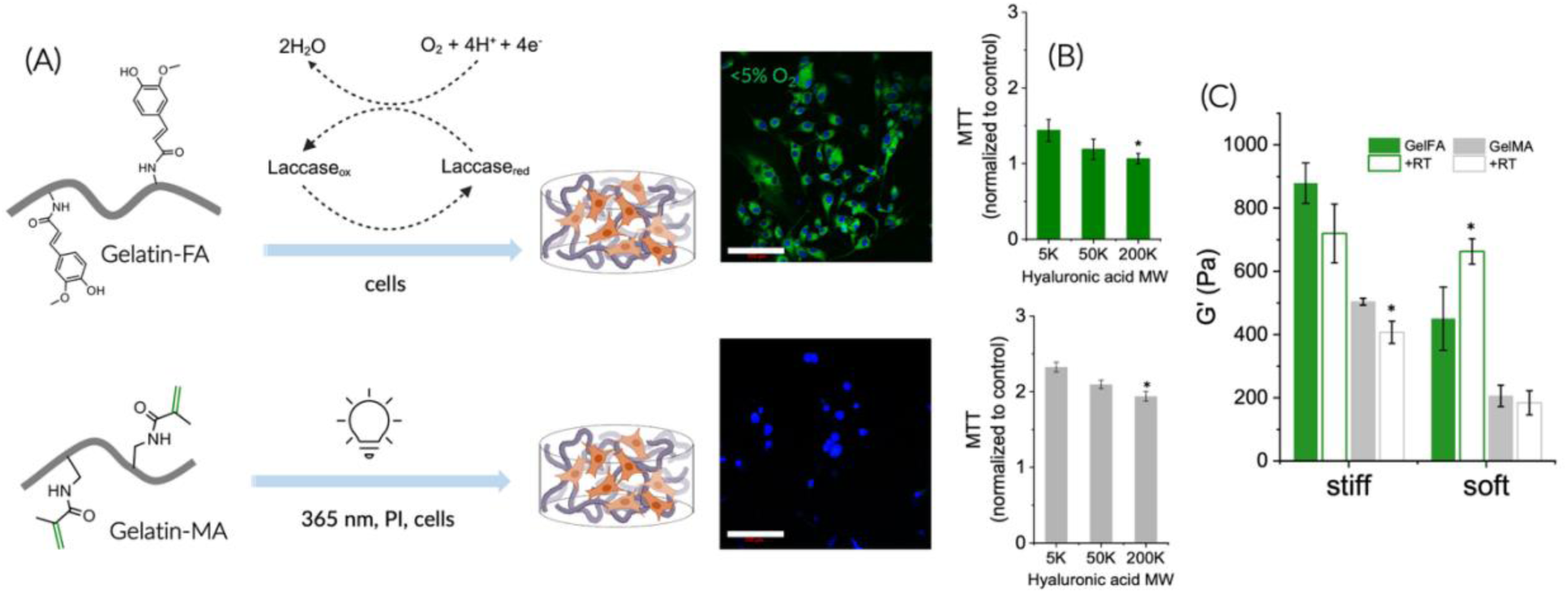
Biomaterial-based environmental control. (A) Gelatin hydrogels are fabricated by using laccase- or UV-initiated reactions to obtain hypoxic (top) and normoxic (bottom) platforms, respectively. Image-iT™ green hypoxia reagent fluorescence is used to determine hypoxia within hydrogels in U87 at less than 5% oxygen, nuclei are shown in blue (DAPI). (B) MTT assay determines the metabolic activity of U87 cells within gelatin hydrogels under hypoxia (top) and normoxia (bottom) when exposed to HA fragments with different molecular weights. (C) Storage moduli (G’) of hydrogels with different stiffnesses before and after radiation therapy. *p<0.05, compared to 5K HA (B) and pre-RT (C).

Increasing the anticancer effects of tumor therapy requires understanding the underlying mechanism for matrix stiffness to promote tumor survival. Post-radiation effects on tissue architecture may involve fibrosis, and vascular damage, that leads to decrease oxygen diffusion and the formation of stiffer and denser fibrous tissue (31). We have found that ionizing radiation-induced changes in matrix stiffness can trigger changes in cell activity. The most significant difference can be seen in the soft matrices, where hypoxic hydrogels show stiffening compared to normoxic scaffolds (unchanged), suggesting cell-induced contraction or remodeling (**Figure 1C**). Mechanical properties of the hydrogels post-radiation indicate an increase of storage modulus for soft GelFA-hypoxic hydrogels, potentially due to increased crosslinking of unreacted ferulic acid groups as a result of additional radical formation (32). For stiffer hydrogels, storage modulus (G’) decreases post-radiation, suggesting potential degradation of gelatin bonds. At low oxygen concentration during irradiation, it has been reported that degradation of gelatin does not occur. On the contrary, increased crosslinking, especially in highly functionalized samples has been widely reported (33). The capacity for these models to predict radiation induced fibrotic processes, involving excessive deposition of extracellular matrix components like hyaluronic acid (HA), paves the way for more effective treatment strategies, with reduced recurrence rate, as reported for some immunotherapeutic approaches (34).

### Extracellular matrix stiffness and oxygen availability influence radiation-induced DNA damage

While studies of the effect of mechanical properties on chemoresistance are extensive, more investigation is necessary to evaluate radio-resistance. Some studies in lung cancer show a more effective radiation-induced DNA damage in stiffer matrices due to chromatin remodeling (35). However, other studies suggest that stiffer environments lead to chromatin compaction, which can protect cells from radiation-induced DNA damage (36). We used GelMA and GelFA hydrogels with adjusted mechanical properties, that contained radiosensitive GBM6 glioma cells (37). Samples were irradiated (single dose of 8 Gy) and tested 1h post-radiation. We found that after cells are irradiated, softer matrices are more likely to promote cellular radiosensitivity: 85% of cells in soft matrices developed γH2AX foci, as compared to 12% in stiffer matrices (6.7 ± 0.6 kPa). We also observed that hypoxic environments reduce the number of double-strand breaks (DSBs) in DNA after radiation (Figure 2). Together, these data suggest that while radiation induces increased DNA damage in softer matrices, with almost ten times increased γH2AX positive cells versus stiffer matrices, damage is significantly reduced and the effect of matrix stiffness far less pronounced for GBM cells in hypoxic environments.

**Figure 2.**
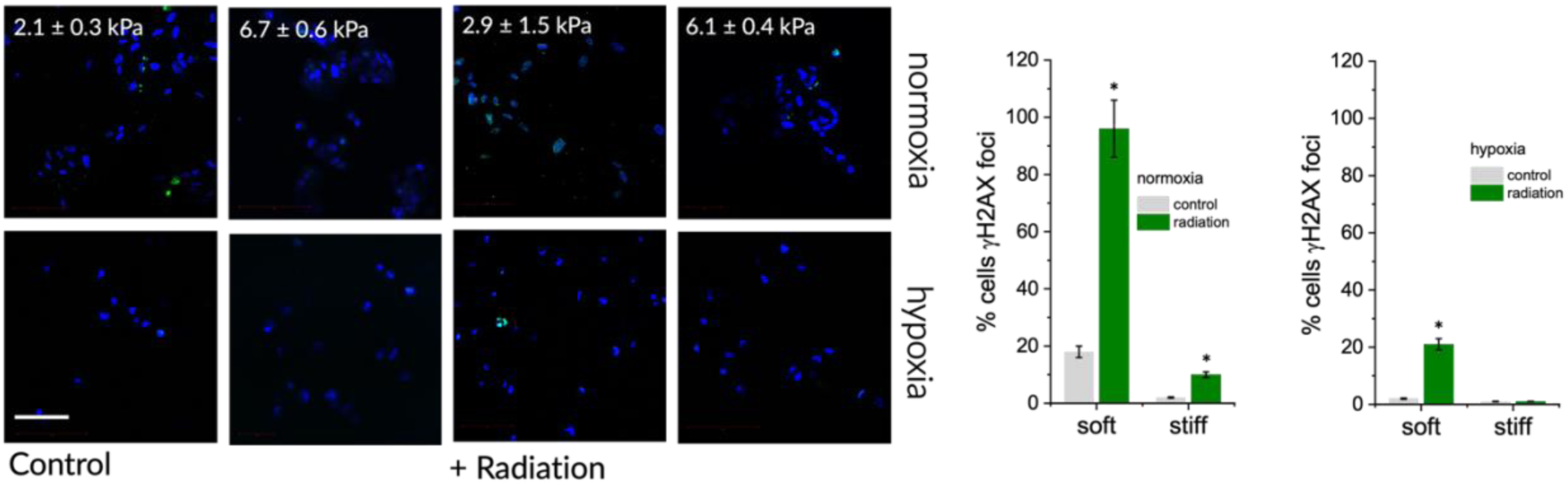
Oxygen concentration and matrix stiffness influence DNA damage after RT. DNA breaks are detected after RT (8 Gy) by using immunofluorescence, γH2AX foci (green) are tested in soft and stiff matrices. The percentage of cells with foci is displayed on right, with soft and normoxic platforms showing the highest concentration. 300 cells were counted per condition. *p<0.05, compared to control.

Hypoxia is a well described factor that influences radio-resistance, but its manner of emergence is also important to consider. The presence of areas of high oxygen consumption in the GBM microenvironment can leads to hypoxia and the inhibition of radical species that produce long-term DNA damage (38,39). Here, we used a ferulic acid mediated crosslinking scheme that consumed oxygen during hydrogel gelation, allowing us to evaluate the role of microenvironmental hypoxia rather than proliferation driven changes in oxygen availability, on GBM radioresistance.

### Hypoxia induces post-radiation cell metabolism changes

Abnormalities of cell metabolism in cancer can lead to predominant aerobic glycolysis and the adaptability of cancer cells to changes in oxygen concentration. Aerobic glycolysis leads to overproduction and release of lactate supporting tumor progression and resistance to treatment (40). Because of these protumor effects, glycolysis and lactate production are potential targets for increasing the efficacy of RT (41). The tumor periphery is characterized by pockets of transient hypoxia due to high oxygen consumption from cancer cells, which can lead to poor RT outcomes (42). We assessed mitochondrial respiration via a Seahorse Cell Mito Stress Test (Agilent) using Agilent Seahorse XFe96 extracellular flux analyzer and a hypoxia chamber to evaluate the oxygen consumption of GBM cells within 3D hydrogel tumor tissues (Figure 3A-B). Notably, GBM cells increase glycolysis in the presence of brain-mimetic HA and hypoxia, we show here the combination of hyaluronic acid metabolites and hypoxia alters GBM metabolism (ATP production, oxygen consumption). While radiotherapy shifts GBM cells from oxidative phosphorylation to glycolysis, the shift is attenuated by the presence of hyaluronan in normoxic conditions. These findings are consistent with prior work that showed mitochondrial respiration increases after radiation, possibly as a result of radiation-induced increase in mitochondrial mass (43). Radiation treatment is also known to increase glycolysis and ATP production rate at acute time points (1h). Interestingly, oxygen consumption significantly increases in a hypoxic environment after radiation, most likely related to reoxygenation processes (44).

**Figure 3.**
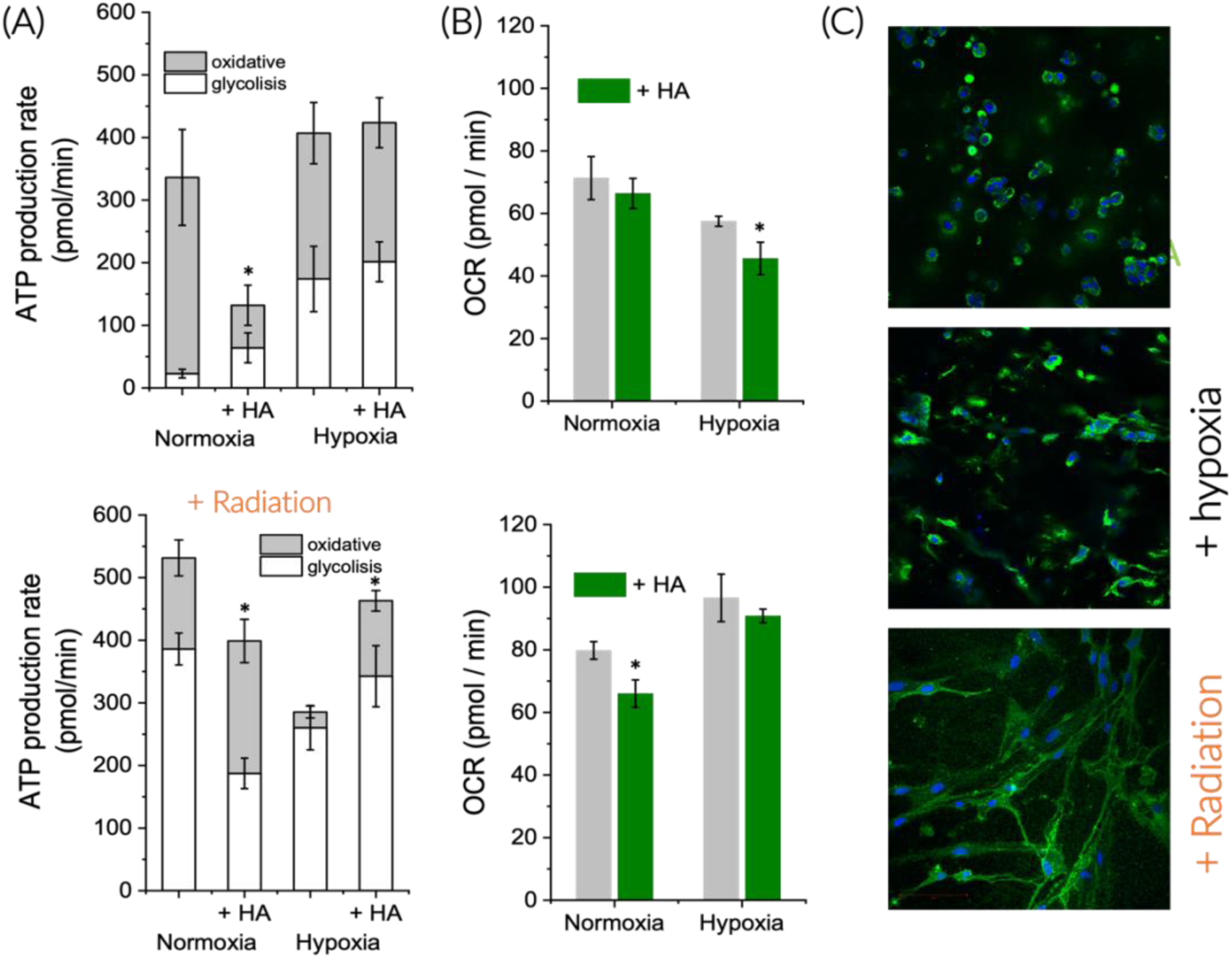
Radiation therapy and hyaluronan influence immediate mitochondrial metabolism in glioma cells. (A) ATP production rate and (B) oxygen consumption of GBM6 cells under hypoxia and when exposed to hyaluronan, using Seahorse (Agilent technologies). (C) HA is imaged inside hydrogels that contain GBM6 glioma cells after RT and under hypoxia. HABP (green) is used to detect the HA with immunofluorescence, nuclei (DAPI, blue). *p<0.05, compared to HA-free samples.

Hyaluronic acid (HA) is the main component of the brain extracellular matrix, and products of its metabolism have been associated with proliferation and invasion. However, it is unclear how overproduction of HA affects metabolism independently of oxygen availability. We have previously described that RT treatment increases HA synthase-2 (HAS2) expression and HA production (29). Increased HA production has also been associated with post-radiation increase of NF-ĸB in vivo (45,46). We show here that cells produce HA after irradiation (Figure 3C) and together with metabolic studies, it indicates that glioma cells can increase glycolytic and oxidative metabolism in response to microenvironmental HA, suggesting a potential positive feedback loop.

Anaerobic metabolism can also modify mitochondrial activity to preserve ATP production independent of oxygen concentration. This effect has been described before as a response to increased extracellular acidosis, leading to alteration of mitochondrial morphology under hypoxic stress (37). Radiation has been associated to changes in the metabolism of GBM in vitro and in vivo, switching into accumulation of lipids (12). We observe that GBM cells immediately respond to 8 Gy RT dose, increasing ATP and glycolysis, and that hyaluronan has a significant role in this response tempering this effect in hypoxia (Figure 3A). Therefore, the modulation of components of the TME may sensitize cells to radiation therapy by decreasing glycolysis.

### Radiation dose rate impacts DNA damage in glioma cells

The detection of γ-H2AX foci is the standard method to quantify DNA double-strand break (DSB) induction and repair (47). Here, we investigated the induction of γ-H2AX foci across a range of patient-derived xenograft GBM lines (37), with each specimen irradiated after day 3 in culture in the GelMA hydrogel. GBM specimens received a single dose of 8 Gy at either a conventional or low dose-rate, with γ-H2AX foci imaged 1h post exposure. Low dose rate (LDR) has been suggested as an alternative to conventional rate to inhibit tumor growth while sparing healthy tissue via intrafraction repair processes (48). Figure 4 shows the characteristics DS breaks induced by radiation, with DNA damage assessed by imaging and quantifying nuclear spots (foci) labeled with fluorescent γ-H2AX DSB repair protein.

**Figure 4.**
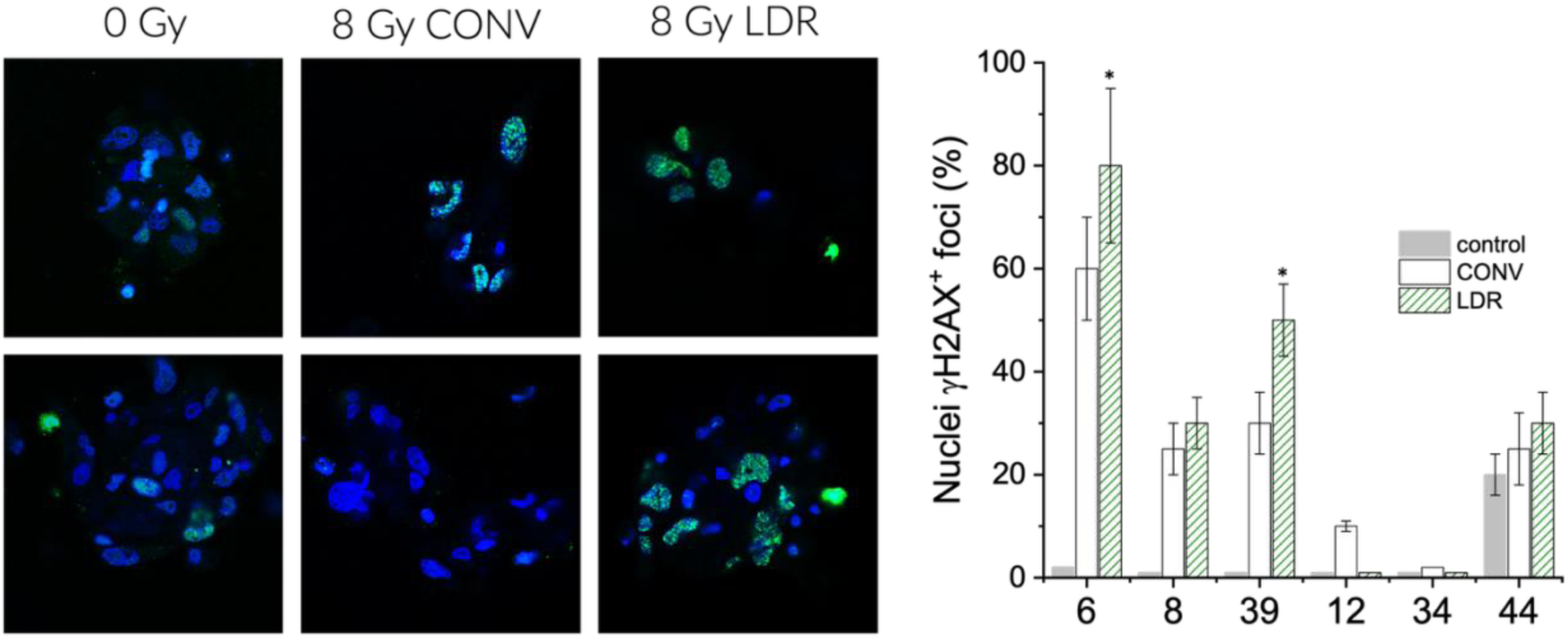
Low dose rate radiation increases DNA damage in radiosensitive GBM cells with EGFR vIII mutations. DNA breaks are detected after RT (8 Gy) by using immunofluorescence, γH2AX foci (green) are tested in cell-laden gelatin platforms. The percentage of cells with foci is displayed on the right. 300 cells were counted per condition. *p<0.05, compared to conventional dose.

Alternative radiation dose rates (ultra-high and low) have the potential to reduce the extent of neurocognitive toxicity associated with brain radiotherapy (49). We observe that the number of γH2AX foci positive cells increase in low dose rate treatments in GBM6, and GBM39, both cell types are EGFRvIII mutant and are characterized as radiosensitive during in vivo experiments (37). Low dose rate advantages lie in the differential DNA repair mechanisms of tumor and healthy cells. Of all the GBM subtypes tested, GBM12 and GBM34 show the highest radioresistance. GBM8 and GBM44 (both p53 wild type tumors) show some radiation sensitivity but no significant difference between conventional and low dose rates. The tumor suppressor protein p53 is essential in DNA repair and becomes relevant in response to radiation-induced damage. Our results support the idea that the combination of EGFR vIII mutation and p53 status can be biomarkers predictive of radiation responsiveness to low dose rates. This platform can serve as a useful tool to assess radiosensitivity before treatment.

### Radiation-induced damage in healthy tissue in combination with antitumor effectiveness

The role of TME in shaping post-radiation metabolic rewiring of cancer cells to induce radio-resistance remains unclear. Here, we used a hydrogel model to assess the degree to which co-cultured neurons may provide a protective effect to GBM during radiation therapy. This approach allows for paracrine communication between different cell populations embedded in 3D biomaterial-based platforms (50–52). Here, we compare the effect of neural cells on GBM cell response for both a well-established GBM cell line (U87) as well as one of the patient-derived xenograft lines that showed strong radio sensitivity (GBM6) (Figure 5A). Interestingly, the presence of neural cells influences glioma response to radiation therapy independent of the 3D culture platform and the cell line. Notably, the presence of neuronal culture attenuates the decrease in metabolic activity post-radiation, regardless of cell type and radiation rate, suggesting a radio-protective effect on GBM cancer cells (Figure 5B).

**Figure 5.**
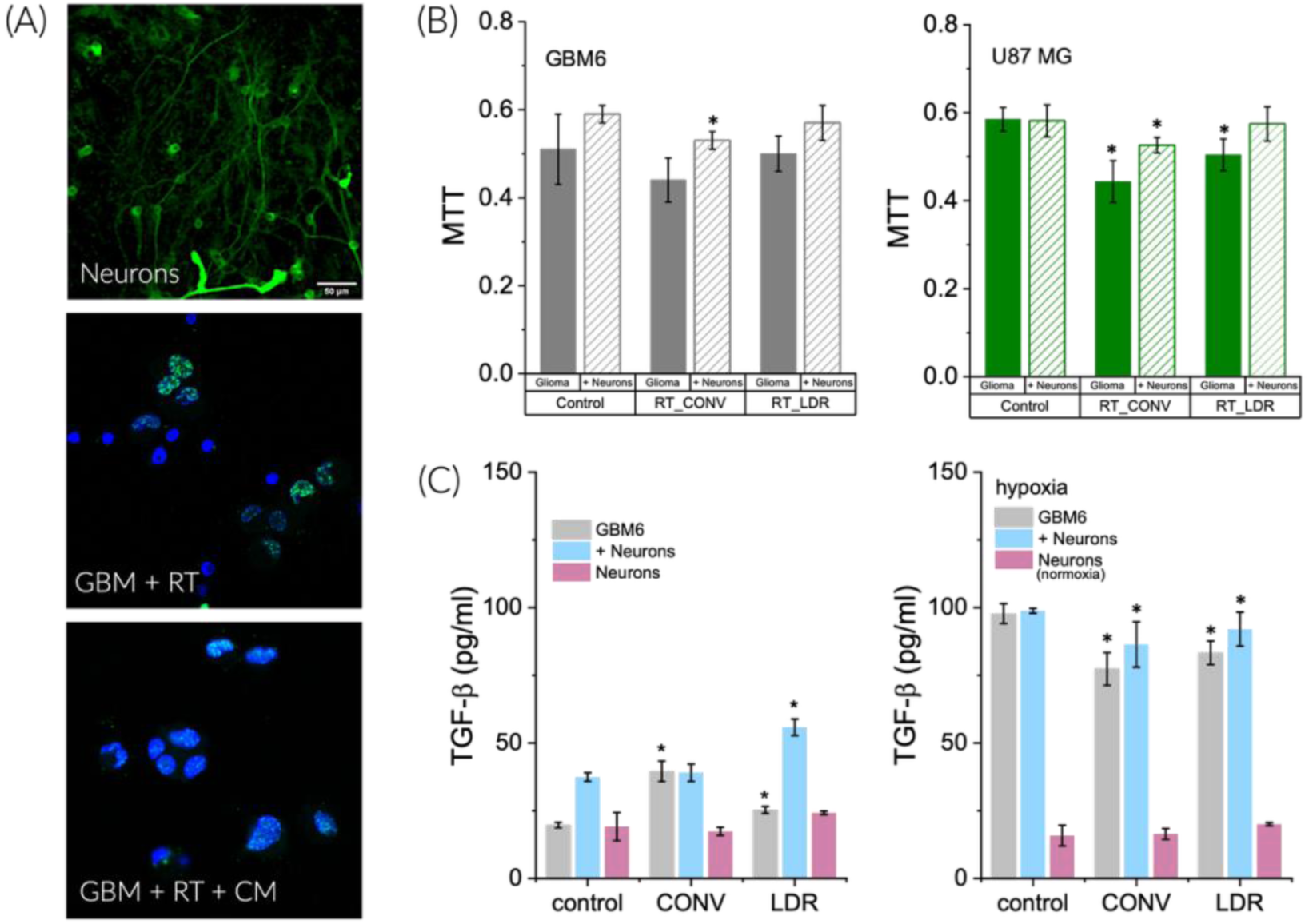
Neurons influence GBM proliferation after RT. (A) Neurons (βIII tubulin, green) and GBM cells (DAPI blue, γ-H2AX foci, green) in 3D hydrogels before and after RT, and in the presence of neuronal conditioned media (CM). (B) MTT assay is used to evaluate the metabolic activity of co-cultured cells (neurons and GBM cells) before and after RT (CONV and LDR). (C) In those cultures, TGFβ was detected with ELISA from the media samples. *p<0.05, compared to control.

The transforming growth factor-beta (TGF-β) signaling pathway is a critical mediator of the response to radiation. TGF-β is expressed in response to ionizing radiation, and cellular distribution in the brain is a predictor of neurotoxicity. TGF-β is recognized to be essential in the development of radiation-induced fibrosis and immunosuppression, with increased TGF-β levels related to synaptic dysfunction and cognitive impairment (53). While many factors affect the radiation toxicity, TGF-β production has been linked to radiation dose and also related to resistance and tissue fibrosis (54). The use of low-dose rate radiotherapy involves a conventional radiotherapy dose but at a rate chosen to lie between the repair thresholds of tumor cells and normal tissues, with the goal of selectively triggering repair only in normal tissues (8,55). This response may be related to the differential expression of TGF-β in normal tissues when exposed to low versus conventional radiation rates. Our data show that GBM6 cells secrete significantly increased TGF-β in response to the presence of neurons, with even increased levels of TGF-β in response to low rate versus conventional radiation but only in normoxic conditions (Figure 5C). Interestingly, secretion of TGF-β is high for GBM6 cells in hypoxia regardless of RT dose rate or presence of neurons. These data manifest the importance of cell-cell interactions in considering GBM radiotherapy, and further that distinctive characteristics of the neuro-glioma unit may shape therapeutic efficacy. Engineered ex vivo tissue models provide an alternative approach to examine complex multicellular interactions that affect radio-resistance in the GBM tumor microenvironment.

### Real-time in-vitro biophysical assessment of glioblastoma cells during radiation

We developed a functional experimental platform integrated within a controlled hypoxic and biomechanically regulated environment that enables quantitative evaluation of glioma cells and neuronal networks during radiation exposure. To achieve real-time, non-perturbative monitoring of radiation response, we employed Spatial Light Interference Microscopy (SLIM), a quantitative phase imaging technique capable of measuring nanoscale cellular structure and dynamics in live cells via interferometry. SLIM enables continuous, label-free monitoring of biophysical changes in GBM cells during and after radiotherapy. Time-resolved cellular responses to fractionated radiation have remained difficult to capture due to the lack of experimental platforms that combine live-cell microscopy with synchronous irradiation. SLIM overcomes this limitation by providing quantitative optical path-length maps with sensitivity comparable to atomic force microscopy, while offering substantially higher acquisition speeds (28).

By interfacing a miniaturized X-ray source directly with a quantitative phase imaging system, we eliminate a key barrier in time-resolved radiobiology, enabling measurements across timescales ranging from milliseconds to days. To capture acute cellular responses to radiation, we performed multimodal SLIM imaging on U87 glioblastoma monolayers, combining quantitative dry-mass measurements with mitochondrial (Mitotracker) and endocytic membranal activity (FM1-43) fluorescence. The nanoscale sensitivity of SLIM to optical path-length fluctuations enables uninterrupted, real-time measurement of nuclear and perinuclear (“ring”) dry-mass dynamics during irradiation, consistent with prior quantitative phase studies of intracellular dry-mass dynamics in living cells (56). A mini-X-ray source (50 kVp, 80 μA) was integrated directly into the SLIM optical path, allowing cells to be imaged immediately before, during, and after RT. Representative SLIM images (Figure 6A) show U87 cell division and neurite extension within neuron–glioma co-cultures. Individual cells, nuclei, and neurite structures were segmented from fluorescence images using Cellpose, a generalist deep-learning–based segmentation framework (57). In these co-cultures, neurite segmentation is observed and serves as a morphological indicator of neuronal injury and impending neuronal death. The analysis of mitochondrial fluorescence revealed pronounced perinuclear mitochondrial clustering (Figure 6B**–D**). This perinuclear mitochondrial accumulation increased during the initial minutes of RT, peaked near the end of treatment, and subsequently declined toward baseline levels, while nuclear-associated mitochondrial signal remained comparatively stable. Corresponding z-score analyses (Figure 6D) confirm that this redistribution is spatially localized to the perinuclear region. This behavior is associated to ROS induced stress and with microtubule depolymerization that ends up causing mitochondrial retrograde trafficking toward the nucleus (58–60). These mitochondrial redistributions temporally coincide with a transient increase in perinuclear dry mass detected by SLIM.

**Figure 6.**
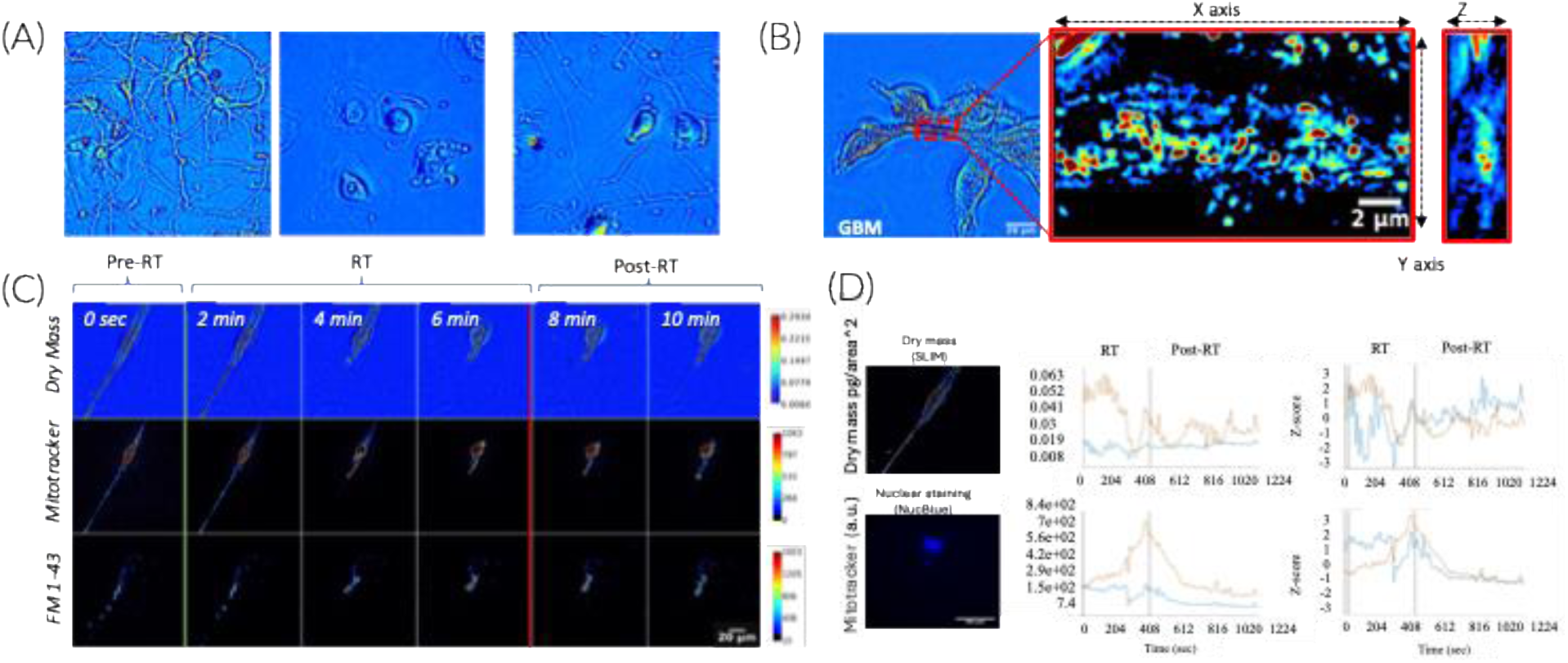
Multimodal SLIM captures rapid dry-mass and mitochondrial redistribution in glioblastoma monolayer cell cultures under acute radiation stress. Label-free imaging: (A) Cortical neurons, glioma cells and the co-culture of both cell types imaged with SLIM and (B) 3D tomographic acquisition. (C) Multimodal acquisition and real time radiation treatment of GBM cells. This montage shows snapshots every 5 frames. (D) Representative multimodal analysis of plasma membrane and mitochondrial dynamics. It is possible to quantify the mean square displacement of single cells and clusters, time-resolved cellular morphology as well as cellular and nuclear dry-mass of incubated GBM-U87 cells during two 20 Gy fractions over 48hours.

Together, these observations are consistent with compensatory cellular responses to acute oxidative stress, including mitochondrial perinuclear clustering, membrane fragmentation, and blebbing (supplementary videos). Perinuclear mitochondrial clustering represents an active, stress-regulated mitochondrial redistribution that is required for hypoxia- and heat-stress–responsive signaling, rather than a byproduct of mitochondrial damage (58,59). These results indicate that radiation induces highly dynamic, spatially compartmentalized changes in dry mass and mitochondrial activity, reflecting an acute, ROS-driven biophysical shock that unfolds over the first several hundred seconds of irradiation. Importantly, these SLIM experiments define the intrinsic timescale of GBM responses in 2D liquid monolayer cultures, where reactive oxygen species and oxygen diffuse freely, enabling direct visualization of the immediate biophysical phase of the radiation response, including rapid dry-mass redistribution and transient perinuclear mitochondrial bursts. In contrast, within 3D hydrogel systems, physiological constraints such as restricted oxygen diffusion and extracellular matrix–dependent mechanical signaling modulate mitochondrial dynamics and slow the progression of molecular responses. As a result, the same oxidative insult observed in 2D should unfolds more gradually in 3D environments, leading to downstream responses that more closely approximate physiological and therapeutic conditions.

## Conclusions

This work describes a hydrogel approach to evaluate glioblastoma cells response to radiotherapy in the context of brain-mimetic hyaluronic acid and hypoxia as well paracrine signaling with neurons. We found that in a subset of patient-derived xenograft specimens, low dose rate radiation induced increased DNA damage compared to standard dose rate. Alternative radiation dose rates have the potential to reduce the extent of neurocognitive toxicity associated with brain radiotherapy. We also observe that the length of neurite outgrowth and number of branches decrease in standard dose rate irradiated samples, providing additional evidence supporting future efforts using low-dose rate irradiation. Our results suggest that combinatorial therapies that control ECM secretion and inflammation may improve the efficacy of RT in glioblastoma. Translating these insights into clinical and therapeutic interventions may guide the design of safer and more effective cancer treatments.

## Acknowledgements

We acknowledge the following institutes for access to their facilities and services: the Roy J. Carver Biotechnology Center, the Tumor Engineering and Phenotyping Core at the Cancer Center at Illinois, and the Carl R Woese Institute for Genomic Biology. Research reported in this publication was supported by The Elsa U. Pardee Foundation (SPH), the National Cancer Institute of the National Institutes of Health under Award Numbers R01 CA256481 (BACH), the Cancer Center at Illinois Planning Grant (SPH) and the William H. Donner Professorship at Mayo Clinic (JNS). The content is solely the responsibility of the authors and does not necessarily represent the official views of the NIH. This work is supported by the National Science Foundation (CHE-2420683 for Y. L.). We also thank the Robert A. Welch Foundation (Grant F-0020) for its support of the Lu group research program at the University of Texas at Austin. The authors are also grateful for additional funding provided by the Department of Chemical & Biomolecular Engineering, the Department of Chemistry, and the Cancer Center at Illinois at the University of Illinois Urbana-Champaign. Figures were created in BioRender.com and OriginLab.

